# scmap - A tool for unsupervised projection of single cell RNA-seq data

**DOI:** 10.1101/150292

**Authors:** Vladimir Yu Kiselev, Andrew Yiu, Martin Hemberg

## Abstract

Single-cell RNA-seq (scRNA-seq) is widely used to investigate the composition of complex tissues^1–9^ since the technology allows researchers to define cell-types using unsupervised clustering of the transcriptome^8,10^. However, due to differences in experimental methods and computational analyses, it is often challenging to directly compare the cells identified in two different experiments. Here, we present scmap (http://bioconductor.org/packages/scmap), a method for projecting cells from a scRNA-seq experiment onto the cell-types or individual cells identified in other experiments (the application can be run for free, without restrictions, from http://www.hemberg-lab.cloud/scmap).

## Main text

As more and more scRNA-seq datasets become available, carrying out comparisons between them is key. A central application is to compare datasets of similar biological origin collected by different labs to ensure that the annotation and the analysis is consistent. Moreover, as very large references, e.g. the Human Cell Atlas (HCA)^11^, become available, an important application will be to project cells from a new sample (e.g. from a disease tissue) onto the reference to characterize differences in composition, or to detect new cell-types (Fig. 1a). Conceptually, such projections are similar to the popular BLAST^12^ method, which makes it possible to quickly find the closest match in a database for a newly identified nucleotide or amino acid sequence.

**Figure 1.**
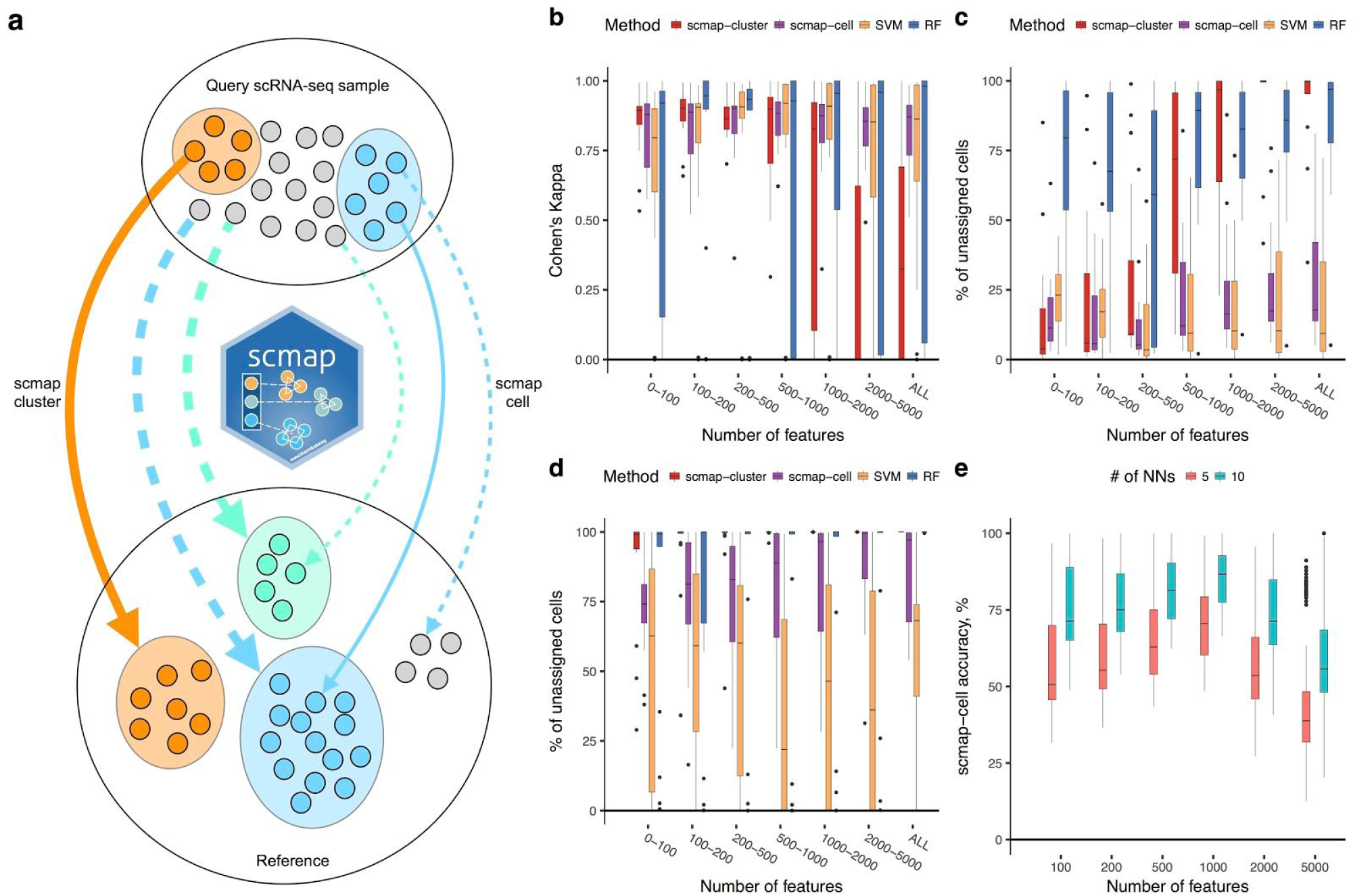
scmap use-cases and performance. (**a**) scmap can be used to compare two different samples by mapping individual cells from a query sample either to cell-types in the reference (scmap-cluster, thick lines) or to individual cells in a reference (scmap-cell, thin lines). The comparison can be carried out either when both samples have been annotated (full lines) or when only one of them is annotated (dashed lines). (**b**) Cohen’s κ values and (**c**) percentage of unassigned cells for positive controls. The plots are based on datasets listed in Table S2 (projections are performed in both directions). Dropout-based feature selection is used for all three methods (see Methods). scmap-cell is run once for each pair of datasets. (**d**) Percentage of unassigned cells in negative controls. The plot is based on datasets listed in Table S3 and projections are performed in both directions. Dropout-based feature selection is used everywhere (see Methods). scmap-cell is run once for each pair of datasets. (**e**) scmap-cell was used to search for nearest neighbours and the accuracy shows how often the true nearest neighbour was found amongst the five or ten nearest cells. The plots are based on datasets listed in Table S1 except Shekhar and Macosko (projections are performed in both directions). Dropout-based feature selection is used (see Methods). scmap-cell is run 100 times for each dataset.

Projecting a new cell, *c*, onto a reference dataset, amounts to identifying which cluster or cell *c* is most similar to, i.e. the nearest neighbor. We represent each cluster by its centroid, i.e. a vector of the median value of the expression of each gene, and we measure the similarity between *c* and each cluster centroid or cell. Searching for the nearest cluster can be done exhaustively since the number of clusters is typically much smaller than the number of cells in the reference. To speed up the search for the nearest cell, we carry out an approximate nearest neighbor (ANN) search using a product quantizer^13^. Moreover, instead of using all genes when calculating the similarity, we use unsupervised feature selection to include only the genes that are most relevant for the underlying biological differences which allows us to overcome batch effects^14^.

We investigate three different strategies for feature selection: random selection, highly variable genes (HVGs)^15^ and genes with a higher number of dropouts than expected (M3Drop)^14^. To increase speed, we modified the M3Drop method and instead of fitting a Michaelis-Menten model to the log expression-dropout relation, we fit a linear model (Methods, Fig. S1a). For the number of features, we used the top 100, 200, 500, 1000, 2000, 5000, or all genes. Similarities were calculated using the cosine similarity, Pearson and Spearman correlations. This has the advantage of being insensitive to differences in scale between datasets as the similarities are restricted to the interval [−1, 1]. To make the cluster assignments more robust, we required that at least two of the three similarities were in agreement, and that their value exceeded.7 for at least one of them. If these criteria are not met, then *c* is labelled as “unassigned” to indicate that it does not correspond to any cell-type present in the reference. For the ANN search, which we refer to as scmap-cell, we carry out a form of k-nearest neighbour classification with only the cosine similarity. The nearest three neighbours are found and we require that they have the same cell-type and that the highest similarity among them to be >.5 for the cell-type to be assigned.

To validate the projections, we considered 17 different datasets^1–9,16–22^ from mouse and human, collected and processed in different ways (Table S1). First, we evaluated different feature selection methods by carrying out a self-projection experiment where each dataset is mapped onto itself. We used 70% of the cells from the original sample for the reference and the remaining 30% were projected, with clusters as defined by the original authors. To quantify the accuracy of the mapping, we use Cohen’s κ^23^ which is a normalized index of agreement between sets of labels that accounts for the frequency of each label. A value of 1 indicates that the projection assignments were in complete agreement with the original labels, whereas 0 indicates that the projection assignment was no better than random guessing. We find that the dropout-based method for feature selection has the best performance, and somewhat surprisingly we also find that random selection is better than HVG (Fig. S1b). Furthermore, the dropout-based method performs consistently well when the number of features selected is in the range of [100, 1000]. The dropout-based method performs better than HVG because it selects genes that are either absent or present in each cluster, and these genes provide a more reliable signal for separating groups of cells^14^. As a comparison, we also considered two commonly used supervised methods for assigning labels to new samples, a random forests classifier (RF) and a support vector machine (SVM). These classifiers were trained on the reference and then applied to the held out cells as before. For the self-projection experiment, we find that both RF and SVM perform slightly better than both scmap-cluster and scmap-cell for all three feature selection methods.

As a positive control, we considered seven pairs of datasets (Table S2) that we expected to correspond well based on their origin. The positive controls represent a more realistic use-case since these comparisons include systemic differences between the reference and projection samples, i.e. batch effects. For example, for three of the pairs, one of the datasets was collected using a full-length protocol and the other was collected using a UMI based protocol. The results showed that despite the substantial differences between the protocols^16,24,25^, both scmap-cluster and scmap-cell on average achieve κ>.75 and assignment rates >75% when the number of features used was between 100 and 500 (Fig. 1b, c, S2a, S3). Even though RF achieves the highest κ of the three approaches it also has the lowest assignment rate (< 50%) indicating that it achieves a high specificity at the cost of low sensitivity. An important feature of scmap is that it is robust to gene dropouts since both the centroids and the nearest neighbour relations are unaffected by the increased frequency of zeros (Fig. S4).

As a negative control, we projected datasets with an altogether different origin from the reference (e.g. mouse retina onto mouse pancreas, Table S3). Reassuringly, we found that both scmap versions categorized >90% of the cells as unassigned when the number of features used was >100 (Fig. 1d). Notably, SVM has a much smaller fraction of unassigned cells than RF and scmap, indicating that it is too lenient in assigning matches. Comparing the evidence across the self-projection experiments, the positive and negative controls, we conclude that scmap with 500 features provides the best performance by balancing high sensitivity and specificity with a low false-positive rate.

We evaluated the scmap-cell by asking how often it was able to identify the true nearest neighbor, as defined by calculating the nearest neighbor exactly, amongst one of the five or ten nearest cells. For 15 of the 17 datasets used earlier, scmap has an average accuracy of 64% or 80%, respectively (Fig. 1e). For the two Drop-seq datasets, scmap-cell performed well in identifying the correct cluster, yet it only achieved an accuracy of ~20% for identifying the nearest neighbor. We hypothesize that deeper sequencing is required for scmap-cell to be able to reliably identify nearest neighbors. The ANN search is most useful for differentiation trajectories where the cells are typically thought to best be represented as a continuum rather than discrete clusters^26^. We evaluated the ANN feature of scmap for trajectories for mouse myoblast differentiation^27^, mouse ES differentiation^28^ and mouse fibroblast to neuron reprogramming^29^. We again found that scmap can correctly identify the nearest neighbor in 76%, 91%, and 94% of the cases (Fig. 2a).

**Figure 2.**
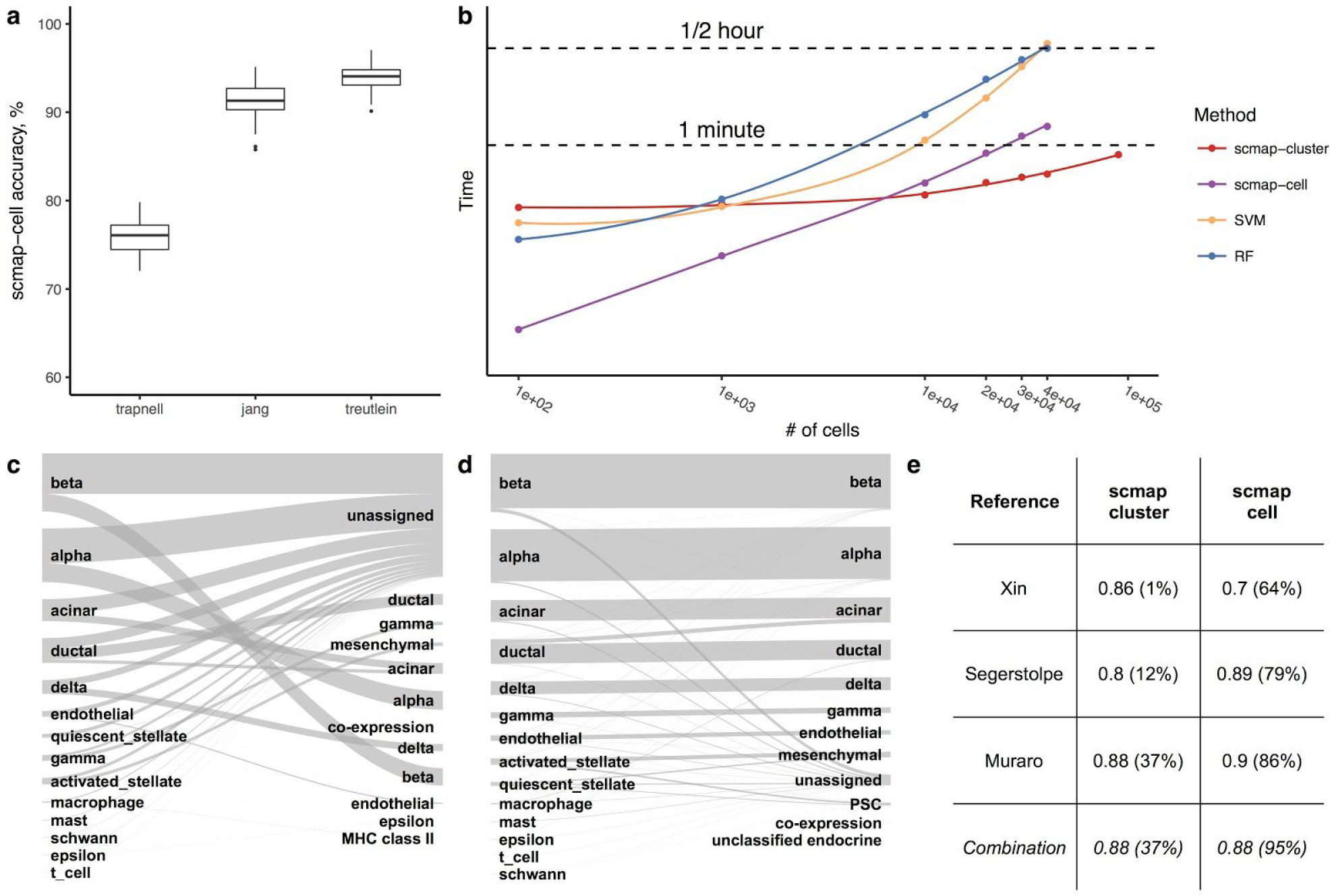
scmap for combined references. (**a**) scmap-cell accuracy for three datasets^27–29^ with differentiation trajectories showing how often the true nearest neighbour was found amongst the ten nearest cells (1000 dropout-selected features were used for projections and scmap-cell was run 100 times for each dataset). (**b**) CPU run times of creating Reference (for scmap) and training classifiers (for SVM and RF). The x-axis represents a number of cells in the reference dataset. For all methods 1,000 features were used. All methods were run on a MacBook Pro laptop (Mid 2014) with 2.8 GHz Intel Core i7 processor, 16 GB 1600 MHz DDR3 of RAM. For 10^5^ cells, scmap-cell failed due to lack of memory. Points are actual data, solid lines are “loess” fit to the points with span = 1 (see ggplot2 documentation). (**c**) Results of scmap-cluster projection of the Baron^4^ dataset to Muraro^2^, Segerstolpe^1^ and Xin^3^ dataset using a combination strategy (Sankey diagram) and (**d**) scmap-cell projection (Sankey diagram). (**e**) Results of scmap-cluster and scmap-cell projections of the Baron^4^ dataset to three human pancreas datasets (Reference) and results of the *Combination* projection.

An important feature of scmap is that it is very fast. It takes only around twenty seconds to select features and to calculate the centroids for 40,000 cells for scmap-cluster, for scmap-cell it takes less than one minute to create the index, whereas it takes almost thirty minutes to train using RF or SVM (Fig. 2b). For all four methods the time to project the new cells is negligible, which means they are very fast with a pre-computed reference. Since the complexity scales with the number of clusters in the reference, rather than the number of cells, scmap-cluster will be applicable to very large datasets as the index is ~5000 times smaller than the original expression matrix. The scmap-cell index is ~500-fold smaller than the original expression matrix (Table S1).

Large references, including the HCA, will be an agglomeration of datasets collected by different groups. Merging different scRNA-seq datasets remains an open problem^30–32^, but the results from our study suggest that samples with similar origin are largely consistent^33^ (Fig. 1c). Instead of correcting for batches and merging, one can create a composite reference and compare the new cells to each dataset separately. When there are multiple datasets in the reference, scmap reports the best match for each dataset. Thus, if a cell shows a high degree of similarity to clusters with similar annotations from different datasets, that will increase the confidence of the mapping. To illustrate the mapping to multiple datasets, we considered the pancreas dataset by Baron *et al^4^* since it had the most unassigned cells when projected to the other pancreas datasets. Combining all projections (Methods) we were able to reduce the fraction of unassigned cells from 99% (Xin^3^) and 88% (Segerstolpe^1^) to 63% while not making κ worse. Interestingly, for this example the performance of scmap-cell was better than scmap-cluster (Fig. 2c-e). Since the reference used by scmap is modular and can be extended without re-calculating the features or centroids for the datasets that have been processed previously, the strategy of not merging datasets is well suited for large references that are expected to grow over a long period of time.

We have implemented scmap as an R-package, and it is part of Bioconductor to facilitate incorporation into bioinformatic workflows. Since scmap is integrated with scater^34^, it is easy to combine with many other popular computational scRNA-seq methods. Moreover, we have made scmap available via the web (http://www.hemberg-lab.cloud/scmap), allowing users to either upload their own reference, or to use a reference collection of datasets from this paper for which the features and centroids have been pre-calculated (Methods).

Due to differences in experimental conditions, comparing scRNA-seq datasets remains challenging. However, for researchers to be able to take advantage of large references, e.g. the HCA, fast, robust and accurate methods for merging^35,36^ and projecting cells across datasets are required. To the best of our knowledge, scmap is the first widely applicable projection method since it can identify both the best matching cell-type as well as individual cell in the reference. We have demonstrated that scmap can be used to compare samples of similar origin collected by different groups, as well as for comparing cells to a large reference composed of multiple datasets.

## Methods

### Datasets

All datasets and cell type annotations were downloaded from their public accessions. The datasets were converted into Bioconductor’s SingleCellExperiment (http://bioconductor.org/packages/SingleCellExperiment) class objects (details are available on our dataset website: https://hemberg-lab.github.io/scRNA.seq.datasets). In the Segerstolpe^1^ dataset cells labeled as “not applicable” were removed since it is unclear how to interpret this label and what it should be matched to in the other datasets. In the Xin^3^ dataset cells labeled as “alpha.contaminated”, “beta.contaminated”, “gamma.contaminated” and “delta.contaminated” were removed since they likely correspond to cells of lower quality. In the following datasets similar cell types were merged together:
All data and scripts
- In the Deng^17^ dataset *zygote* and *early2cell* were merged into *zygote* cell type, *mid2cell* and *late2cell* were merged into *2cell* cell type, and *earlyblast*, *midblast* and *lateblast* were merged into *blast* cell type.
- All bipolar cell types of the Shekhar^9^ dataset were merged into *bipolar* cell type.
- In the Yan^21^ dataset *oocyte* and *zygote* cell types were merged into *zygote* cell type.

### Feature selection

To select informative features we used a method conceptually similar to M3Drop^14^ to relate the mean expression (*E*) and the dropout rate (*D*). We used a linear model to capture the relationship log(*E*) and log(*D*), and after fitting a linear model using the lm() command in R, important features were selected as the top *N* residuals of the linear model (Fig. S1a). The features are only selected from the reference dataset, and those of them absent or zero in the projection dataset are are further excluded before running scmap. All three feature selection methods are described in Supplementary Note 1.

### Reference centroid

In scmap-cluster each cell type in the reference dataset is represented by its centroid, i.e. the median value of gene expression for each feature gene across all cells in that cell type.

### Approximate nearest neighbor search using product quantizer

scmap-cell performs fast approximate k-nearest neighbour search using product quantization^13^. The original algorithm, built around the Euclidean distance, was adapted to incorporate the cosine distance, which helps to protect against batch effects and scaling inconsistencies between datasets. The product quantizer creates a compressed index where every cell in the reference is identified with a set of sub-centroids found via k-means clustering based on a subset of the features. By concatenating the sub-centroids, a close approximation to the original expression vector is obtained. When searching the reference for the nearest neighbours to a query cell, the approximations provided by the sub-centroids are used instead of the individual cells in the reference. Since the number of centroids can be made much smaller than the original number of cells in the dataset, the method provides a substantial reduction in both computation time and storage requirements compared to exact search.

### Projection dataset

Projection of a dataset to a reference dataset is performed by calculating similarities between each cell and all centroids of the reference dataset, using only the common selected features. Three similarity measures are used: Pearson, Spearman and cosine. The cell is then assigned to the cell type which correspond to the highest similarity value. However, scmap-cluster requires that at least two similarity measures agree with each other, otherwise the cell is marked as “unassigned”. Additionally, if the maximum similarity value across all three similarities is below a similarity threshold (default is.7), then the cell is also marked as “unassigned”. Since only the cosine similarity measure was calculated for scmap-cell, the default threshold of.5 was used, and the nearest three neighbours are required to agree on the cell-type for it to be assigned. Positive and negative control plots corresponding to Figs. 1c-e for different values of the similarity/probability (see *SVM and RF*) threshold (.5,.6,.8 and.9) are shown in Fig. S2.

### SVM and RF

The scmap projection algorithm was benchmarked against support vector machines^37^ (with a linear kernel) and random forests^38^ (with 50 trees) classifiers from the R packages e1071 and randomForest. The classifiers were trained on all cells of the reference dataset and a cell type of each cell in the projection dataset was predicted by the classifiers. Additionally, a threshold (default value of.7) was applied on the probabilities of assignment: if the probability was less than the threshold the cell was marked as “unassigned”.

### Sensitivity to sequencing depth and dropouts

The dropout rate in the positive control datasets (Table S2) was artificially increased by randomly setting 10%, 30% and 50% of the non-zero expression values to zero (Fig. S4a,b). scmap was run 100 times for each box.

### Projection based on multiple datasets

When the reference contains multiple datasets collected from similar samples by different groups in addition to all similarities, for each cell scmap also reports a top cell type match based on the highest value of similarities across all reference cell types. A similarity threshold of.7 is also applied in this case.

### scmap on the Cloud

An example of a cloud version of scmap is available at http://www.hemberg-lab.cloud/scmap. Instructions of how to set it up on a user’s personal web cloud environment are available on our github page: https://github.com/hemberg-lab/scmap-shiny. An extended tutorial on how to use scmap can be found in Supplementary Note 2.

### Figures

All data and scripts used to generate figures in this paper are available at https://github.com/hemberg-lab/scmap-paper-figures.

## Acknowledgements

We would like to thank Tallulah Andrews, Kedar N Natarajan, Guillermo Parada, Michael Schaub, Michael Stubbington, Valentine Svensson, Jennifer Westoby and Florian Wünnermann for helpful discussions, feedback on the manuscript and for testing the cloud implementation of scmap.

## Funding

Amazon Web Services (AWS) Cloud provided credits for running the scmap server for one year. VYK and MH were both supported by core funding to the Wellcome Trust Sanger Institute provided by the Wellcome Trust.

## Conflicts of interest

None.

## Author contributions

M.H. conceived the study and supervised the research; V.Y.K., A.Y. and M.H. contributed to the computational framework; V.Y.K. and M.H. wrote the manuscript.

